# Different subtypes of EGFR exon19 mutation can affect prognosis of patients with non-small cell lung adenocarcinoma

**DOI:** 10.1101/374710

**Authors:** Yingying Tian, Jiuzhou Zhao, Pengfei Ren, Bo Wang, Chengzhi Zhao, Chao Shi, Bing Wei, Jie Ma, Yongjun Guo

**Affiliations:** Department of Molecular Pathology, Henan Cancer Hospital, The Affiliated Cancer Hospital, Zhengzhou University, Zhengzhou, China; School of Pharmaceutical Sciences, Zhengzhou University, Zhengzhou, China

**Keywords:** EGFR exon19, mutation subtype, non-small cell lung adenocarcinoma, prognosis

## Abstract

**Aims:** In this study, we determined whether different subtypes of epidermal growth factor receptor (EGFR) exon19 mutation are associated with the therapeutic effect of EGFR-tyrosine kinase inhibitors (TKIs) on advanced non-small cell lung adenocarcinoma.

**Methods:** A total of 122 patients with stage III or IV non-small cell lung adenocarcinoma were retrospectively reviewed. Clinical characteristics of these patients, including progression-free survival (PFS) outcome for EGFR-TKI treatment, were analyzed.

**Results:** According to the mutation pattern, we classified the in-frame deletions detected on EGFR Exon19 into three different types: codon deletion (CD), with a deletion of one or more original codons; codon substitution and skipping (CSS), with a deletion of one or two nucleotides but the residues could be translated into a new amino acid without changing following amino acid sequence; CD or CSS plus single nucleotide variant (SNV) (CD/CSS+SNV), exclude CD or CSS, there’s another SNV nearby the deletion region. The clinical characteristics of three groups were analyzed and as a result, no significant difference was found. By comparing the average number of missing bases and amino acids of the three mutation subtypes, it could be discovered that the number of missing bases and amino acids of the three mutation subtypes is diverse, and group CSS> group CD> group CD/CSS+SNV. Finally, survival analysis was performed between three groups of patients. The median PFS of group CD, group CSS and group CD/CSS+SNV was 11 months, 9 months and 14 months respectively. There was a distinct difference in the PFS between group CSS and group CD/CSS+SNV (P=0.035<0.05), and the PFS of group CD/CSS+SNV was longer.

**Conclusions:** Different mutation subtypes of EGFR exon19 can predict the therapeutic effect of EGFR-TKIs on advanced non-small cell lung adenocarcinoma.

## Introduction

Lung cancer is the leading cause of cancer-related death worldwide[1]. Non-small cell lung cancer (NSCLC) accounts for about 85% of lung cancer[2, 3]. With the emergence of EGFR-TKIs, such as gefitinib, erlotinib, and icotinib, the survival time and life quality of patients with lung cancer have been obviously improved [4-6]. Provided the inconsistent effect of EGFR-TKIs on different patients, some patients received EGFR-TKIs with apparent efficacy, while others did not work. Therefore, it is particularly important to screen out these patients who showing a better response to EGFR-TKIs.

Epidermal growth factor (EGF) and its receptor (EGFR), which is most closely related to lung adenocarcinoma, were discovered by Stanley Cohen of Vanderbilt university in 1986[7]. Human EGFR gene is located on chromosome 7p11.2, which contains 28 exons, sizing about 200kb. EGFR exon 18 ~ 24 encodes tyrosine kinase functional region of receptor. The vast majority of EGFR mutation occur in NSCLC, especially in non-smoking Asian women[8-10]. Compared to patients with negative EGFR mutation, EGFR-mutated patients showed longer PFS (11.5 months)[11]and overall survival(OS) (15.4 months)[12]. The common subtypes of EGFR mutation include exon19 deletion (19del) and point mutation on exon21(L858R), accounting for 33.1% and 40.9% respectively[13]. Multiple randomized clinical trials have demonstrated that EGFR 19del and L858R are highly correlated with sensitivity to EGFR-TKI in NSCLC, especially 19del[14, 15]. However, Patients with EGFR 19 del results different PFS after EGFR TKI treatment, and the reason is still not clear. This retrospective study reviewed the medical records of EGFR exon19 mutant advanced NSCLC patients undergoing EGFR-TKIs treatment, so as to evaluate the association of different subtypes of exon19 mutation with EGFR-TKI efficacy in EGFR-mutant advanced non-small cell lung adenocarcinoma patients.

## Materials and Methods

### Patients

Inclusion criteria were as follows: (1) patients with pathologically confirmed non-small cell lung adenocarcinoma who underwent EGFR mutation screening and TKIs treatment at Henan Cancer Hospital between 2015 and 2017;(2) patients who merely harbored EGFR exon19 mutation and did not receive other treatment before targeted therapy. A total of 122 patients were collected between 2015 and 2017 in Henan Cancer Hospital. EGFR exon19 mutation was detected in tumor tissues by the method of Next Generation Sequencing (NGS). All subjects were administered with gefitinib(n=76), erlotinib (n = 10), or icotinib (n = 36), until disease progression or intolerance to adverse events. Since the above three EGFR-TKIs have similar therapeutic effect[16], this study did not take into account the different impact of targeted drugs such as gefitinib, erlotinib and icotinib on patients’ PFS.

### Detection of EGFR exon19 mutation by NGS

gDNA from formalin-fixed paraffin-embedded (FFPE) tissues was extracted using a QIAamp^®^ Circulating Nucleic Acid kit (Qiagen, Hilden, Germany), according to the manufacturer’s recommendations. All gDNA samples were first assessed using a NanoDrop-2000(Thermo Scientific, Wilmington, DE, USA), and then qualified by a Qubit^®^ 2.0 Fluorometer (Invitrogen, Carlsbad, CA, USA) using a Qubit^®^ dsDNA HS Assay Kit. gDNA was fragmented to about 200bp with Diagenode SA. Libraries were constructed using a human polygenic mutation detection kit (Burning Rock Dx), which analyzes target regions of 8 genes (EGFR, KRAS, BRAF, ERBB2, ALK, ROS1, RET, and MET). The library size was checked using the Agilent High Sensitivity DNA Kit by the Bioanalyzer 2100 instrument (Agilent Technologies), and library concentration was evaluated with a Qubit^®^ 2.0 Fluorometer using the Agilent High Sensitivity DNA Kit (Life Technologies), following the manufacturer’s instructions. Finally, sequencing was performed on the Illumina^®^ NextSeq™ 550.

### Follow up

All 122 patients were followed up through hospital medical records until disease progression or April 2018. In this study, response was classified by standard Response Evaluation Criteria in Solid Tumors (RECIST). Confirmed by imaging, the total maximum diameter of the target lesion increased by at least 20%, or new lesion appeared, which was defined as disease progression. PFS was defined as the duration from administration of EGFR-TKIs to disease progression. The object of study had a median follow-up period of 8 months (range, 1 to 53 months), with a 100% follow-up seen. During the follow-up period, 82 patients have been observed with progression.

### Statistics

All statistical analyses were conducted using the statistical software SPSS version 16.0. Univariate analyses were performed with a log-rank test. Clinical characteristics of the different groups, including gender, age, smoking history, drinking history, family history and Eastern Cooperative Oncology Group Performance Status (ECOG PS) were compared by χ2 test or Fisher exact test, and the survival was estimated with the Kaplan-Meier method. All P values were 2-sided and P<0.05 was considered statistically significant.

## Results

### Clinical characteristics of patients with EGFR exon19 mutation

A total of 122 lung adenocarcinoma patients with EGFR exon19 mutation were enrolled in this study. The participants included 55 men and 67 women, and had a median age of 57 years (range, 27 to 81 years); 100 cases at ages of 67 years or less, 22 cases aged over 67 years. Of the total study subjects, 30 cases were identified with history of smoking or alcohol drinking while 92 cases were neither smokers nor drinkers; 25 patients were discovered with family history of cancer and 97 patients were not. The ECOG score of 108 cases was 0, 14 cases were 1. According to the TNM staging system for NSCLC[17], 14 patients were identified as staged III, and 108 patients were staged IV. 122 patients had not received any other therapies before EGFR-TKIs treatment. The clinical characteristics of 122 patients were analyzed by univariate analysis, it was found that mentioned clinical data did not affect the survival of patients with EGFR exon19 mutation (all P values > 0.05) (Table 1).

**Table 1.** Univariate analysis of 122 patients with EGFR exon19 mutation.

| Characteristics | No. of patients<br>N=122(%) | Median<br>PFS(months,95%CI) | *P* |
| --- | --- | --- | --- |
| **Gender** |  |  | 0.612 |
| Male | 55(45.1) | 13(12.089-13.911) |  |
| Female | 67(54.9) | 10(8.481-11.519) |  |
| **Age(years)** |  |  | 0.12 |
| $\leq 67$ | 100(82) | 12(10.318-13.682) | |
| $> 67$ | 22(18) | 9(4.361-13.639) | |
| **Smoking** |  |  | 0.708 |
| Yes | 30(24.6) | 12(7.387-16.613) |  |
| No | 92(75.4) | 11(9.376-12.624) |  |
| **Drinking** |  |  | 0.894 |
| Yes | 30(24.6) | 13(11.961-14.039) |  |
| No | 92(75.4) | 11(9.384-12.616) |  |
| **Family history** |  |  | 0.332 |
| Yes | 25(20.5) | 12(8.342-15.658) |  |
| No | 97(79.5) | 11(9.173-12.827) |  |
| **ECOG PS** |  |  | 0.132 |
| 0 | 108(88.5) | 12(10.42-13.58) |  |
| 1 | 14(11.5) | 10(3.781-16.219) |  |

PFS, progression-free survival; CI, confidence interval; ECOG PS, eastern cooperative oncology group performance status.

### Distribution of EGFR exon19 mutation subtypes in NSCLC

There were 16 mutation subtypes in this study, including c.2235_2249del, c.2236_2250del, c.2239_2247del, c.2240_2254del, c.2237_2251del, c.2237_2254del, c.2240_2257del, c.2235_2251delins, c.2235_2252delins, c.2236_2248delins, c.2236_2255delins, c.2237_2255delins, c.2237_2257delins, c.2239_2248delins, c.2239_2251delins, c.2252_2276delins, respectively (Table 2). In this study, we divided all patients into three groups based on EGFR exon19 mutation pattern: group CD, group CSS and group CD/CSS+SNV (Fig 1). The clinical data of three groups were analyzed by χ2 test. It was discovered that there was no significant difference in the clinical data between three groups (Table 3).

**Fig 1.**
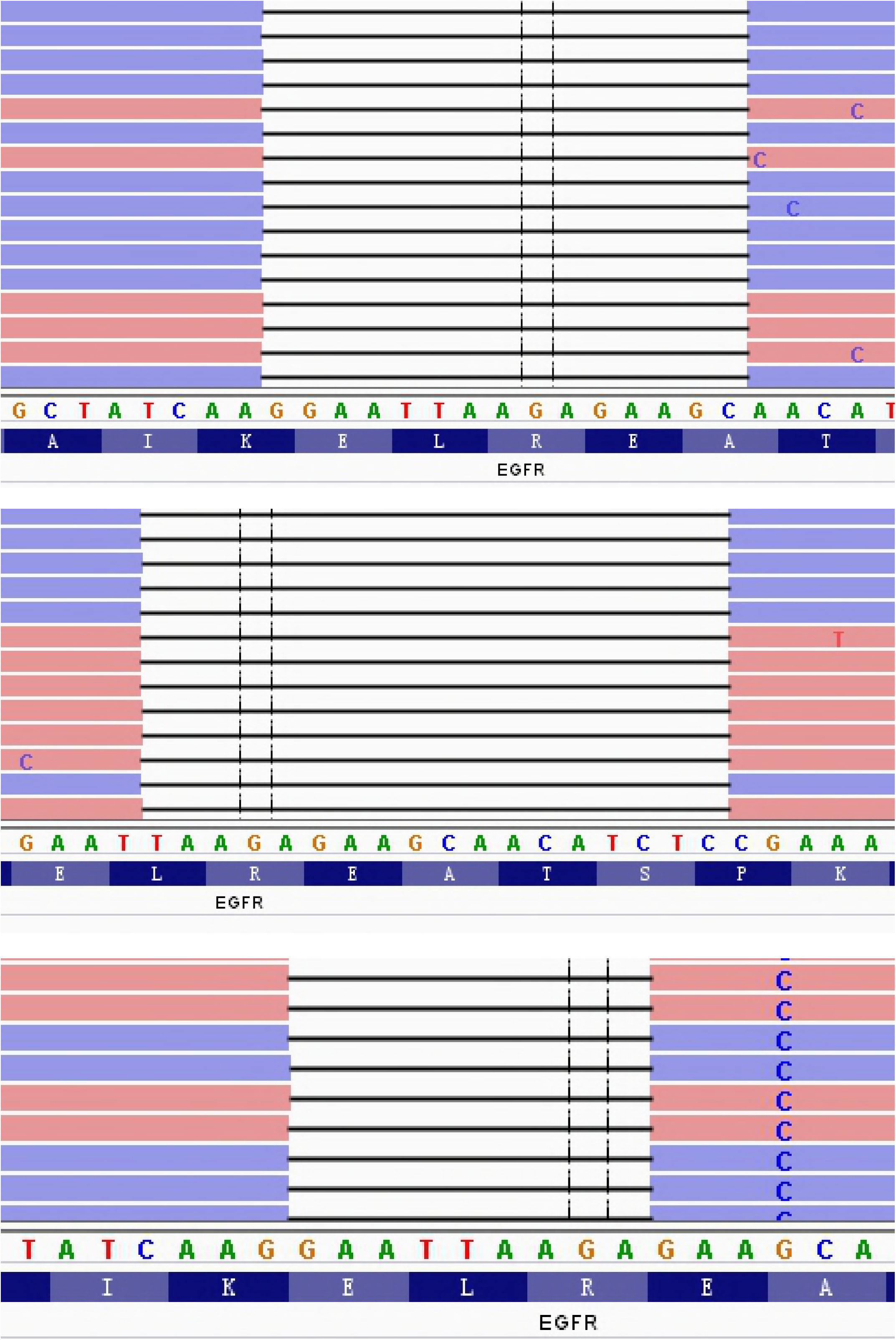
Three mutation subtypes of EGFR exon19. (a) CD, codon deletion; (b) CSS, codon substitution and skipping; (c) CD/CSS+SNV, CD or CSS plus SNV

**Table 2.** Mutation subtypes of 122 EGFR exon19 mutation.

| Base change | Amino acids change | N (%) |
| --- | --- | --- |
| c.2235_2249del | p.Glu746_Ala750del | 49 (40.2) |
| c.2236_2250del | p.Glu746_Ala750del | 32 (26.2) |
| c.2239_2247del | p.Leu747_Glu749del | 1 (0.8) |
| c.2240_2254del | p.Leu747_Thr751del | 8 (6.6) |
| c.2237_2251del | p.Glu746_Thr751delins | 4 (3.3) |
| c.2237_2254del | p.Glu746_Ser752delins | 1 (0.8) |
| c.2240_2257del | p.Leu747_Pro753delins | 10 (8.2) |
| c.2235_2251delins | p.Glu746_Thr751delins | 1 (0.8) |
| c.2235_2252delins | p.Glu746_Thr751delins | 1 (0.8) |
| c.2236_2248delins | p.Glu746_Ala750delins | 1 (0.8 ) |
| c.2236_2255delins | p.Glu746_Ser752delins | 1 (0.8 ) |
| c.2237_2255delins | p.Glu746_Ser752delins | 4 (3.3 ) |
| c.2237_2257delins | p.Glu746_Pro753delins | 1 (0.8 ) |
| c.2239_2248delins | p.Leu747_Ala750delins | 5 (4.2 ) |
| c.2239_2251delins | p.Leu747_Thr751delins | 2 (1.6 ) |
| c.2252_2276delins | p.Thr751_Ile759delins | 1 (0.8 ) |

**Table 3.**
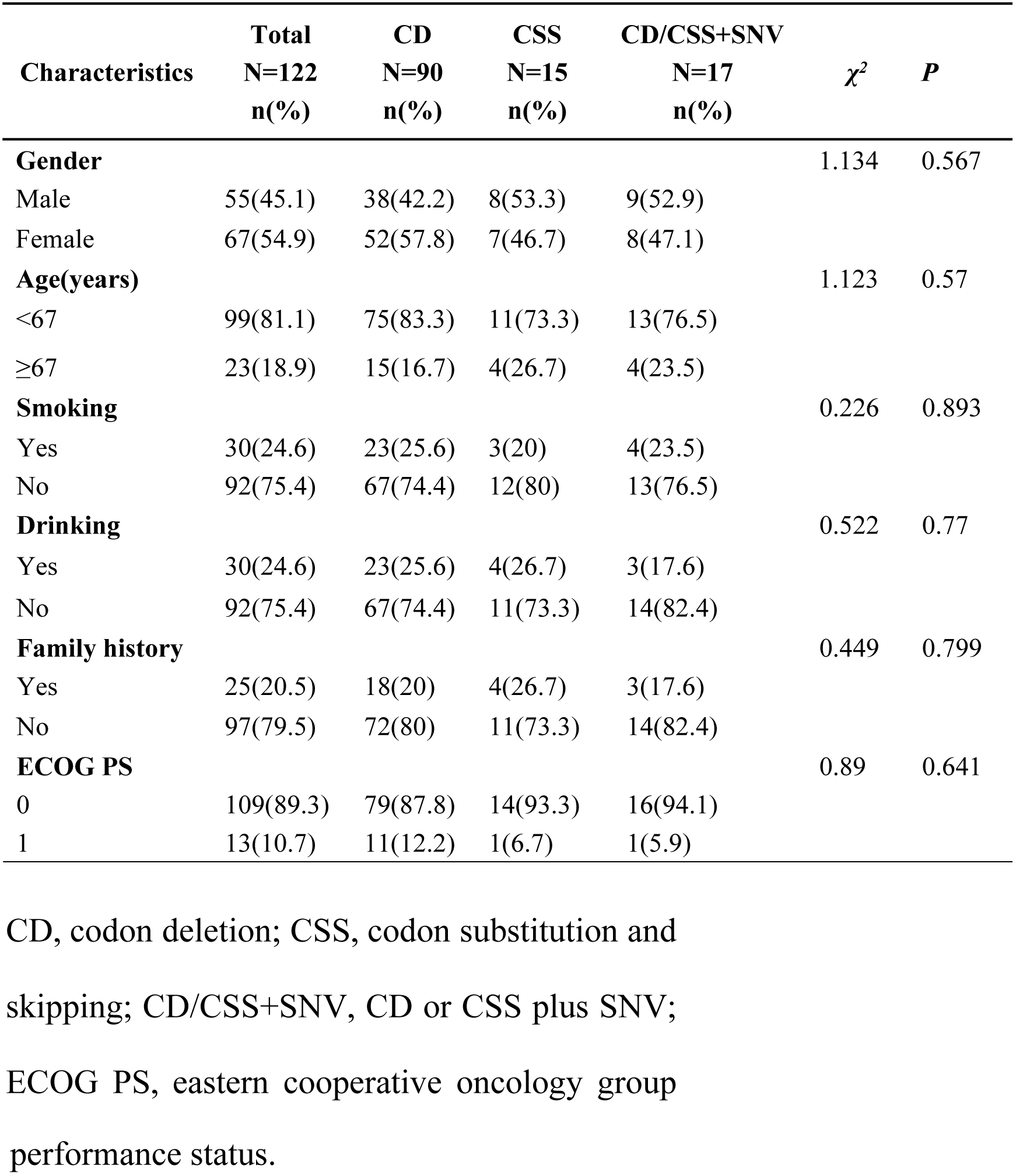
Clinical characteristics of three groups.

| Characteristics | Total<br>N=122<br>n(%) | CD<br>N=90<br>n(%) | CSS<br>N=15<br>n(%) | CD/CSS+SNV<br>N=17<br>n(%) | $\chi^2$ | P |
| --- | --- | --- | --- | --- | --- | --- |
| **Gender** |  |  |  |  | 1.134 | 0.567 |
| Male | 55(45.1) | 38(42.2) | 8(53.3) | 9(52.9) |  |  |
| Female | 67(54.9) | 52(57.8) | 7(46.7) | 8(47.1) |  |  |
| **Age(years)** |  |  |  |  | 1.123 | 0.57 |
| <67 | 99(81.1) | 75(83.3) | 11(73.3) | 13(76.5) |  |  |
| ≥67 | 23(18.9) | 15(16.7) | 4(26.7) | 4(23.5) |  |  |
| **Smoking** |  |  |  |  | 0.226 | 0.893 |
| Yes | 30(24.6) | 23(25.6) | 3(20) | 4(23.5) |  |  |
| No | 92(75.4) | 67(74.4) | 12(80) | 13(76.5) |  |  |
| **Drinking** |  |  |  |  | 0.522 | 0.77 |
| Yes | 30(24.6) | 23(25.6) | 4(26.7) | 3(17.6) |  |  |
| No | 92(75.4) | 67(74.4) | 11(73.3) | 14(82.4) |  |  |
| **Family history** |  |  |  |  | 0.449 | 0.799 |
| Yes | 25(20.5) | 18(20) | 4(26.7) | 3(17.6) |  |  |
| No | 97(79.5) | 72(80) | 11(73.3) | 14(82.4) |  |  |
| **ECOG PS** |  |  |  |  | 0.89 | 0.641 |
| 0 | 109(89.3) | 79(87.8) | 14(93.3) | 16(94.1) |  |  |
| 1 | 13(10.7) | 11(12.2) | 1(6.7) | 1(5.9) |  |  |

CD, codon deletion; CSS, codon substitution and skipping; CD/CSS+SNV, CD or CSS plus SNV; ECOG PS, eastern cooperative oncology group
performance status.

Furthermore, Independent- Samples T Test was conducted for comparing the number of missing bases and missing amino acids of different mutation types. The mean number of missing bases in group CD was 14.93, 17.2 in group CSS and 14.18 in group CD/CSS+SNV. The results showed significant differences in the number of missing bases between group CD and group CD/CSS+SNV, likewise in group CSS and group CD/CSS+SNV (P<0.05). The mean number of missing amino acids in group CD was 4.98, 5.73 in group CSS and 4.76 in group CD/CSS+SNV. Consistently, there were also significant differences in the number of missing amino acids between group CD and group CSS, similarly in group CSS and group CD/CSS+SNV (P<0.05) (Table 4).

**Table 4.** The number of missing bases and amino acids of three groups.

| Characteristics | CD<br>Mean | CSS<br>Mean | *P* | CD<br>Mean | CD/CSS+SNV<br>Mean | *P* | CSS<br>Mean | CD/CSS+SNV<br>Mean | *P* |
| --- | --- | --- | --- | --- | --- | --- | --- | --- | --- |
| Missing base number | 14.93 | 17.2 | < 0.01 | 14.93 | 14.18 | 0.548 | 17.2 | 14.18 | 0.029 |
| Missing amino acid number | 4.98 | 5.73 | <0.01 | 4.98 | 4.76 | 0.572 | 5.73 | 4.76 | 0.022 |

CD, codon deletion; CSS, codon substitution and skipping; CD/CSS+SNV, CD or CSS plus SNV

### Impact of EGFR exon19 mutation subtypes on prognosis of patients

There were 90 patients in group CD, of whom 59 cases had progress and 31 cases had no progress; 15 patients in group CSS, 10 cases of whom had progress, 5 cases of whom did not. Group CD/CSS+SNV included 17 patients, 10 cases of whom had progress, while 7 cases of whom were not. Furthermore, the kaplan-meier survival analysis was performed on three groups. The median PFS in group CD was 11 months, the 95% confidence interval was 9.427-12.573, the median PFS in group CSS was 9 months, the 95% confidence interval was 3.97-14.03, the median PFS in group CD/CSS+SNV was 14 months, and the 95% confidence interval was 9.862-18.138. It was found that there was a significant difference in PFS between group CSS and group CD/CSS+SNV (P<0.05). The median PFS of patients in group CD/CSS+SNV is significantly higher than that in group CD and group CSS (Fig 2), which indicates that different mutation subtypes of EGFR exon19 can influence the therapeutic effect of EGFR-TKIs on advanced non-small cell lung adenocarcinoma.

**Fig 2.**
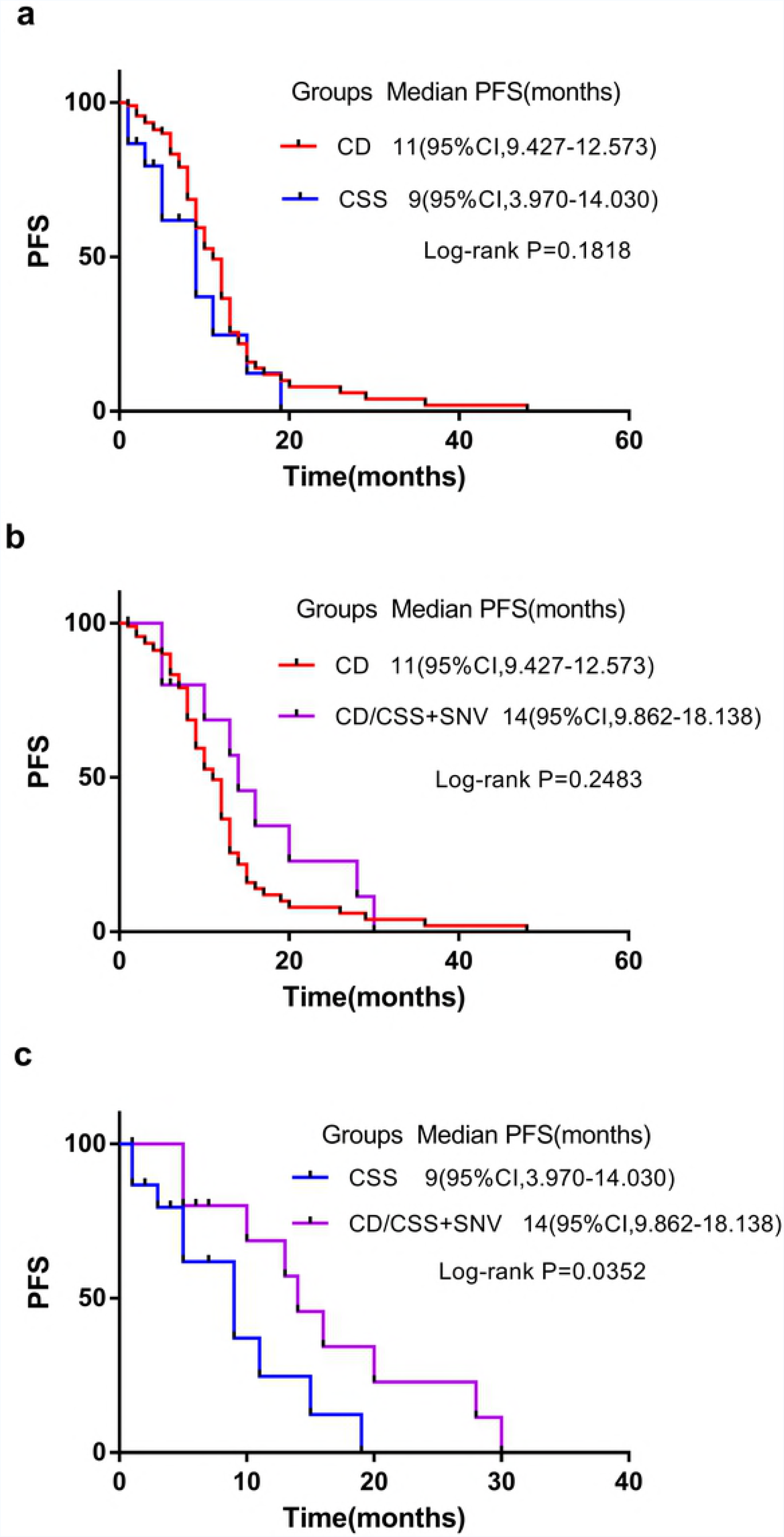
Kaplan-Meier curves of progression-free survival (PFS). (a) CD, codon deletion; (b) CSS, codon substitution and skipping; (c) CD/CSS+SNV, CD or CSS plus SNV

## Discussion

A total of eight genes were detected by NGS in our study, including EGFR, KRAS, BRAF, ERBB2, ALK, ROS1, RET and MET, and it is common-accepted that EGFR gene mutation accounted for the majority. Various randomized clinical trials have emphasized that EGFR 19del and 21L858R mutation in NSCLC are highly sensitive to EGFR-TKIs, especially 19del[18, 19]. Currently, due to the limitation of gene detection technology, no institution has studied the effect of mutation subtypes of EGFR exon19 on the prognosis of disease yet. The molecular pathology department of Henan Cancer Hospital introduced the Next 550 sequencer, which has a sensitivity up to 1/1000 for gene sequencing, providing a useful tool for us to study the mutation subtypes of EGFR exon19. This study is a retrospective analysis of all patients with EGFR exon19 mutation in Henan Cancer Hospital from 2015 to 2017. As far as we know, this is the first study to explore the effect of EGFR exon19 mutation subtypes in terms of the prognosis of disease, aims to guide patients with advanced lung cancer to obtain more efficient treatment.

Wei B[20] implicated that tumor genesis driven by different EGFR mutations were mechanistically different. Zhang C[21] showed that patients with mutation of T790M had poor prognosis. As we all know, PFS and OS for advanced NSCLC patients with an exon 19 mutation was considerably higher than those with other subtypes of EGFR mutations[22, 23]. However, due to the individual imparity in the PFS of patients with EGFR exon19 mutation, we tried to figure out these patients who would have a better response from EGFR-TKIs treatment. According to the different pattern of EGFR exon19 mutation, patients were divided into group CD, group CSS and group CD/CSS+SNV. the survival of three groups were analyzed and found that the median PFS of group CD was 11 months, while group CSS was 9 months, and group CD/CSS+SNV was 14 months. Obviously, group CD/CSS+SNV had a longer PFS. Although the survival curves of group CD and group CD/CSS+SNV, group CD and group CSS were not considerably different(P>0.05), it was not difficult to find that the PFS of most patients in group CD/CSS+SNV is higher than that in group CD, and the PFS of most patients in group CD is higher than that in group CSS. Independent-Samples T Test was conducted for the number of missing bases and missing amino acids of different mutation types. The results showed that the number of missing bases and amino acids in group CD/CSS+SNV was the lowest, and the impact of group CD/CSS+SNV on the genome sequence was minimal, which may be the main reason for the longer PFS in group CD/CSS+SNV. In the future, we will engage on the mechanisms of such significant difference.

In conclusion, our study showed that EGFR-TKIs had the best therapeutic efficacy on the group CD/CSS+SNV, which had the longest PFS in patients with EGFR exon19 mutation. Thus, EGFR exon19 mutation subtypes should be considered when predicting prognosis. However, the response of group CSS to EGFR-TKIs was relatively poor, combination therapy could be considered. This study still has limitations. Firstly, the study results may be skewed by the numbered cases of EGFR exon19 mutation; Secondly, the study results may be also influenced on account of the instability of retrospective analysis. More data would be needed in the future to confirm the findings. To sum up, our study screened out patients with EGFR exon19 mutation who had a better response to EGFR-TKIs, which contributes to guiding the clinical treatment of advanced non-small cell lung adenocarcinoma.

## Supporting information

S1 File. The original data of this study.

## Acknowledgments

We thank Zhizhong Wang and Jun Li for their linguistic assistance during the preparation of this manuscript.

## References

1. Torre LA, Bray F, Siegel RL, Ferlay J, Lortet-Tieulent J, Jemal A. Global cancer statistics[J]. CA Cancer J Clin. 2015, 65(2):87–108.

2. He YY, Zhang XC, Yang JJ, Niu FY, Zeng Z, Yan HH, et al. Prognostic significance of genotype and number of metastatic sites in advanced non-small-cell lung cancer[J]. Clin Lung Cancer. 2014, 15(6):441–7.

3. Chen Z, Fillmore CM, Hammerman PS, Kim CF, Wong KK. Non-small-cell lung cancers: a heterogeneous set of diseases[J]. Nat Rev Cancer. 2014, 14(8):535–46.

4. Kuiper JL, Heideman DA, Wurdinger T, Grunberg K, Groen HJ, Smit EF. Rationale and study design of the IRENE-trial (NVALT-16): a phase II trial to evaluate iressa rechallenge in advanced NSCLC patients with an activating EGFR mutation who responded to an EGFR-TKI used as first-line or previous treatment[J]. Clin Lung Cancer. 2015, 16(1):60–6.

5. Lee JK, Hahn S, Kim DW, Suh KJ, Keam B, Kim TM, et al. Epidermal growth factor receptor tyrosine kinase inhibitors vs conventional chemotherapy in non-small cell lung cancer harboring wild-type epidermal growth factor receptor: a meta-analysis[J]. Jama. 2014, 311(14):1430–7.

6. Sunaga N, Tomizawa Y, Yanagitani N, Iijima H, Kaira K, Shimizu K, et al. Phase II prospective study of the efficacy of gefitinib for the treatment of stage III/IV non-small cell lung cancer with EGFR mutations, irrespective of previous chemotherapy[J]. Lung Cancer. 2007, 56(3):383–9.

7. S C. Epidermal growth factor[J]. Nobel lecture. 1986:1981–90.

8. Zhang X, Chang A. Molecular predictors of EGFR-TKI sensitivity in advanced non-small cell lung cancer[J]. Int J Med Sci. 2008, 5(4):209–17.

9. Tanaka T, Matsuoka M, Sutani A, Gemma A, Maemondo M, Inoue A, et al. Frequency of and variables associated with the EGFR mutation and its subtypes[J]. Int J Cancer. 2010, 126(3):651–5.

10. Helland A, Skaug HM, Kleinberg L, Iversen ML, Rud AK, Fleischer T, et al. EGFR gene alterations in a Norwegian cohort of lung cancer patients selected for surgery[J]. J Thorac Oncol. 2011, 6(5):947–50.

11. Tamura K, Okamoto I, Kashii T, Negoro S, Hirashima T, Kudoh S, et al. Multicentre prospective phase II trial of gefitinib for advanced non-small cell lung cancer with epidermal growth factor receptor mutations: results of the West Japan Thoracic Oncology Group trial (WJTOG0403)[J]. Br J Cancer. 2008, 98(5):907–14.

12. Sutani A, Nagai Y, Udagawa K, Uchida Y, Koyama N, Murayama Y, et al. Gefitinib for non-small-cell lung cancer patients with epidermal growth factor receptor gene mutations screened by peptide nucleic acid-locked nucleic acid PCR clamp[J]. Br J Cancer. 2006, 95(11):1483–9.

13. Zhou J, Song X-B, He H, Zhou Y, Lu X-J, Ying B-W. Prevalence and Clinical Profile of EGFR Mutation In Non-Small-Cell Lung Carcinoma Patients in Southwest China[J]. Asian Pacific Journal of Cancer Prevention. 2016, 17(3):965–71.

14. Zhang Y, Sheng J, Kang S, Fang W, Yan Y, Hu Z, et al. Patients with exon 19 deletion were associated with longer progression-free survival compared to those with L858R mutation after first-line EGFR-TKIs for advanced non-small cell lung cancer: a meta-analysis[J]. PLoS One. 2014, 9(9):e107161.

15. Liu JJ, Zhang S, Wu CJ, Ma LX, Liu Y, Li H, et al. Comparison of clinical outcomes of patients with non-small cell lung cancer harboring different types of epidermal growth factor receptor sensitive mutations after first-line EGFR-TKI treatment[J]. Zhonghua Zhong Liu Za Zhi. 2016, 38(3):211–7.

16. Haspinger ER, Agustoni F, Torri V, Gelsomino F, Platania M, Zilembo N, et al. Is there evidence for different effects among EGFR-TKIs? Systematic review and meta-analysis of EGFR tyrosine kinase inhibitors (TKIs) versus chemotherapy as first-line treatment for patients harboring EGFR mutations[J]. Crit Rev Oncol Hematol. 2015, 94(2):213–27.

17. Tsim S, O’Dowd CA, Milroy R, Davidson S. Staging of non-small cell lung cancer (NSCLC): a review[J]. Respir Med. 2010, 104(12):1767–74.

18. Zhou J, Ben S. Comparison of therapeutic effects of EGFR-tyrosine kinase inhibitors on 19Del and L858R mutations in advanced lung adenocarcinoma and effect on cellular immune function[J]. Thorac Cancer. 2018, 9(2):228–33.

19. Wei WE, Mao NQ, Ning SF, Li JL, Liu HZ, Xie T, et al. An Analysis of EGFR Mutations among 1506 Cases of Non-Small Cell Lung Cancer Patients in Guangxi, China[J]. PLoS One. 2016, 11(12):e0168795.

20. Wei B, Ren P, Zhang C, Wang Z, Dong B, Yang K, et al. Characterization of common and rare mutations in EGFR and associated clinicopathological features in a large population of Chinese patients with lung cancer[J]. Pathol Res Pract. 2017, 213(7):749–58.

21. Zhang C, Wei B, Li P, Yang K, Wang Z, Ma J, et al. Prognostic value of plasma EGFR ctDNA in NSCLC patients treated with EGFR-TKIs[J]. PLoS One. 2017, 12(3):e0173524.

22. Wang H, Huang J, Yu X, Han S, Yan X, Sun S, et al. Different efficacy of EGFR tyrosine kinase inhibitors and prognosis in patients with subtypes of EGFR-mutated advanced non-small cell lung cancer: a meta-analysis[J]. J Cancer Res Clin Oncol. 2014, 140(11):1901–9.

23. Choi YW, Jeon SY, Jeong GS, Lee HW, Jeong SH, Kang SY, et al. EGFR Exon 19 Deletion is Associated With Favorable Overall Survival After First-line Gefitinib Therapy in Advanced Non-Small Cell Lung Cancer Patients[J]. Am J Clin Oncol. 2018, 41(4):385–90.

